# Structural and Functional Insights into GGCX-FIX Interaction: Implications for Vitamin K-Dependent Bleeding Disorders

**DOI:** 10.1101/2025.02.18.638829

**Authors:** Kang Liu, Shixin Li, Jiangbo Tong, Nan Jiang, Minwen Hong, Yi Gu, Luju Chen, Dan Liang, Yongchao Jin, Yuan Zhao, Dongmei Hou, Jinlin Huang, Jian-Ke Tie, Zhenyu Hao

**Affiliations:** College of Bioscience and Biotechnology, Yangzhou University, Yangzhou 225009, China; Joint International Research Laboratory of Agriculture and Agri-Product Safety, Ministry of Education of China, Yangzhou, Jiangsu 225009, China; Department of Biology, the University of North Carolina at Chapel Hill, Chapel Hill, North Carolina 27599, USA; Affiliated hospital, Yangzhou University, Yangzhou, Jiangsu 225009, China

## Abstract

Gamma-carboxylation, catalyzed by γ-glutamyl carboxylase (GGCX), is a critical post-translational modification essential for the biological activity of vitamin K-dependent proteins (VKDPs). Mutations in GGCX, depending on their specific location, result in vitamin K-dependent coagulation factor deficiency type 1 (VKCFD1), which encompasses a broad spectrum of clinical manifestations ranging from mild to severe, including bleeding disorders, osteoporosis, and vascular calcification. The limited knowledge of GGCX’s structure and functional regions hinders our understanding of the consequences of GGCX mutations and the treatment for VKCFD1. This study aimed to identify key functional regions of GGCX and their interactions with VKDPs to better elucidate the molecular mechanisms underlying these diverse clinical symptoms. Using AlphaFold 3 and molecular dynamics simulations, we developed a complex binding model of GGCX, FIX, and reduced vitamin K, which revealed critical regions and residues involved in their interaction. Site-directed mutagenesis and cell-based assays further validated the model, confirming that multisite and regional cooperative binding of FIX to GGCX plays a key role in modulating γ-carboxylation efficiency. Additionally, novel residues (I296, M303, M401, M402) were identified as essential for GGCX’s dual enzymatic activities: carboxylation and vitamin K epoxidation. We further demonstrated that the spatial proximity of these active sites supports the hypothesis that GGCX’s carboxylation and vitamin K epoxidation centers are interconnected, ensuring the efficient coupling of these processes. Our GGCX-FIX binding and carboxylation model aligns with known pathogenic GGCX mutations, providing valuable insights into the molecular basis of coagulation disorders caused by GGCX mutants.

## Introduction

Gamma-carboxylation is a critical post-translational modification essential for the biological activity of vitamin K-dependent proteins (VKDPs), which include coagulation factors II (prothrombin), VII, IX, and X, as well as regulatory proteins C, S, and Z. ^1–3^ These proteins are vital for maintaining the balance of blood clotting and anticoagulation.^2^ Additionally, VKDPs such as Matrix Gla Protein (MGP) and Bone Gla Protein (BGP or osteocalcin) play pivotal roles in vascular health and bone metabolism.^4^ The enzyme γ-glutamyl carboxylase (GGCX) catalyzes the carboxylation of specific glutamic acid residues within VKDPs, converting them into γ-carboxyglutamic acid (Gla) residues.^1^ This modification enables VKDPs to bind calcium ions, triggering conformational changes that are essential for their biological activities, including coagulation, bone mineralization, and the inhibition of vascular calcification.^5,6^ Vitamin K acts as a necessary cofactor in this reaction, undergoing a cyclical process of reduction to vitamin K hydroquinone, involvement in γ-carboxylation, and subsequent oxidation to vitamin K epoxide.^7,8^ Efficient recycling of vitamin K is crucial for maintaining GGCX activity.^9^ Pathogenic mutations in the GGCX gene result in vitamin K-dependent coagulation factor deficiency type 1 (VKCFD1), a rare autosomal recessive disorder.^10,11^ While primarily characterized by coagulation abnormalities, the condition also possibly presents with osteoporosis and arterial calcification^12,13^.

Most GGCX pathogenic mutants result in the aberrant substrate binding or abolished enzymatic activity.^2,14^ GGCX recognizes VKDPs by binding to their propeptides, which tether the VKDPs to the enzyme, facilitating proper carboxylation.^15,16^ Reduced affinity between GGCX and VKDPs is associated with a spectrum of bleeding disorders, from mild to severe. For instance, the L394R mutation^17,18^ results in severe bleeding, primarily due to impaired GGCX-VKDP binding. Notably, bleeding symptoms caused by partial GGCX mutations that disrupt its substrate binding can be mitigated with high-dose vitamin K administration.^19^ Similarly, mutations in the propeptide region of coagulation factor IX (e.g., A-10T, A-10D) reduce the GGCX-Coagulation factor IX(FIX) affinity, leading to abnormal carboxylation and symptoms resembling Hemophilia B or warfarin hypersensitivity.^20–22^ However, the precise location of the propeptide binding site on GGCX remains controversial.

In addition to bleeding disorders, some patients with VKCFD1 present with atypical symptoms, such as PEX-like manifestations, which are rare in coagulation disorders.^10^ GGCX mutations, including D153G and R476H/C, differentially affect the carboxylation of coagulation-related and non-hemostatic VKDPs, such as Matrix Gla Protein (MGP).^23,24^ Truncation at R704 results in a significant reduction in BGP carboxylation activity and a moderate decrease in FIX activity, contributing to the onset of VKCFD1^14,25^. Studies have demonstrated that BGP primarily interacts with the C-terminal region of GGCX through its Gla domain.^14,26^ These suggest that GGCX interacts with coagulation factors and non-coagulation VKDPs through distinct binding mechanisms. However, the precise binding model between GGCX and its diverse substrates remains elusive.

Pathogenic mutations in GGCX, such as F299S^24^ and S300F^27^ result in severely diminished or undetectable VKDP function and clinical manifestations resembling VKCFD1 and PEX-like disorders. Further study has confirmed that these mutations are associated with defects in vitamin K epoxidation and carboxylation^14^, and their symptoms cannot be alleviated with vitamin K administration. These findings underscore the importance of understanding GGCX’s active sites to elucidate the mechanisms underlying these effects, facilitating targeted treatment strategies.

GGCX is an integral membrane protein with multiple functional regions that include binding sites for glutamate, propeptides, reduced vitamin K, CO_2_, and the active sites for both carboxylation and vitamin K epoxidation.^1,2^ The interplay between these regions is critical for GGCX’s dual functionality in catalyzing γ-carboxylation and maintaining the vitamin K cycle. Despite extensive research, the structural characterization of these regions and their cooperation in catalysis remains incomplete, leaving the γ-carboxylation mechanism inadequately understood.

To address gaps in the structure-function understanding of GGCX, we constructed a complex binding model incorporating GGCX, FIX, and reduced vitamin K, utilizing AlphaFold 3 and molecular dynamics simulations. This model identified key functional regions and residues critical for GGCX-FIX interaction. Experimental data confirmed that multiple binding sites contribute to the interaction, emphasizing the dynamic nature of γ-carboxylation and the importance of regional cooperation. We further characterized novel active sites involved in both carboxylation and vitamin K epoxidation. Based on these findings, we proposed a model illustrating the coupling of VKDP carboxylation with the vitamin K redox cycle. The constructed GGCX-FIX binding and carboxylation model aligns with known pathogenic mutations in GGCX, validating its reliability and providing insights into the pathophysiology of GGCX-related disorders and informing potential therapeutic strategies.

## Materials and methods

### Materials and cell lines

Materials used include warfarin, trypsin, and Vitamin K1 (MedChemExpress, China), Xfect transfection reagent (Clontech), mammalian expression vectors pBudCE4.1 and pcDNA3.1Hygro(+) (Invitrogen), HEK293 and HEK293T cell lines (ATCC), mouse anti-carboxylated FIX Gla domain antibody (Green Mountain Antibodies), and HRP-conjugated sheep anti-human Protein C IgG and goat anti-mouse IgG (Affinity Biologicals, Jackson ImmunoResearch). All cell culture media and oligonucleotide primers were also sourced from Invitrogen. (San Diego, CA).

### Simulation methods

To forecast the binding modes of GGCX with FIX, we first used AlphaFold 3^28^, a powerful tool for the predication of protein-protein interactions, to obtain the binding complex of GGCX-FIX. Besides, the binding complexes of GGCX-PC and GGCX-BGP were also generated to compare with GGCX-FIX binding mode. Meanwhile, based on FIX-GGCX binding mode, three mutations of FIX propeptide, including F-6R, A-10T and F-16Y, and GGCX mutation H410P were generated using PyMOL^29^ to compare the contributions of GGCX and FIX propeptide in their binding. Based on PC-GGCX binding model, the GGCX mutation of H755P, E757V and F758L were used for comparation to probe the role of GGCX C-terminal in its catalytic activity. The molecular docking was performed to revealed the binding mode of GGCX with Vitamin K using AutoDock Vina^30^ (version 1.1.2). Molecular dynamics (MD) simulations were performed to evaluate the conformational change of FIX propeptide mutations and the stability of all binding complexes, based on Gromacs (version 2020.6)^31^. The binding contribution of amino acid residues within GGCX was calculated using MM/PBSA method^32^. The details for Molecular docking, MD simulations and MM/PBSA calculation were shown in Supplementary files.

### DNA Manipulations and Plasmid Constructions

Mammalian expression vector pcDNA3.1, with the cDNA of FIX, Protein C, FIXgla-PC (PC with its Gla domain exchanged with that of FIX) was used as the cloning and expression vector, as previously described^14^. The BGP was fused with Protein C and cloned into the expression vector. All other chimeric reporter-proteins, with different propeptides and/or Gla domains used in the cell-based functional study were obtained by overlap PCR. The nucleotide sequences of all the constructs were verified by DNA sequencing at Genewiz Inc. (Suzhou, P. R. China).

### Measurement of reporter-protein carboxylation

Carboxylation activity of reporter proteins was assessed in HEK293 cells as previously described.^23,33^ Plasmid DNA containing chimeric reporter protein cDNA was stably expressed in HEK293 cells lacking endogenous GGCX. Wild-type and mutant GGCX were transiently expressed in these cells. After 48 hours in 5 µM Vitamin K1, the medium was collected for ELISA to measure carboxylated proteins using mouse anti-carboxylated FIX Gla domain antibody (capture) and GAFIX-HRP (detection). Data were analyzed using GraphPad Prism 8. For ELISA standards, chimeric proteins were purified from HEK293/VKOR cells.

### Bimolecular fluorescence complementation (BiFC) analysis

The affinity of VKDP reporters for GGCX was evaluated using bimolecular fluorescence complementation (BiFC) analysis, which has been decried before^14^. We split yEm Venus into two fragments and fused them with GGCX or reporter proteins to create complementary pairs. BiFC pairs were co-transfected into GGCX-deficient HEK293 cells. The affinity variation between GGCX and different reporter proteins can be assessed by monitoring fluorescence at Excitation 515nm/Emission 530nm by the microplate reader of Enspire MLD 2300 (PerkinEImer).

### Assessment of GGCX’s epoxidase activity

High-performance liquid chromatography (HPLC) was employed to access the epoxidation activity of GGCX or mutants^33^. Wild-type GGCX or mutant variants are transfected into HEK293 cells lacking endogenous GGCX gene. After 40 hours incubation with warfarin (5 µM), replaced the medium with 20 µM Vitamin K1 for another 6 hours. Collected the cells and determined KO formation from KH_2_ with HPLC, providing a quantitative measure of epoxidation activity.

## Results

### Multisite and regional cooperative binding of FIX to GGCX modulates γ-Carboxylation efficiency

To elucidate the interaction of VKDPs with GGCX, we employed AlphaFold 3 to develop a detailed binding model of FIX in complex with GGCX. The model predicts that FIX (green-purple-red-blue) interact with the GGCX framework (gold) (**Figure 1A**). The spatial arrangement of these residues offers crucial insights into potential contact regions.

**Figure 1.**
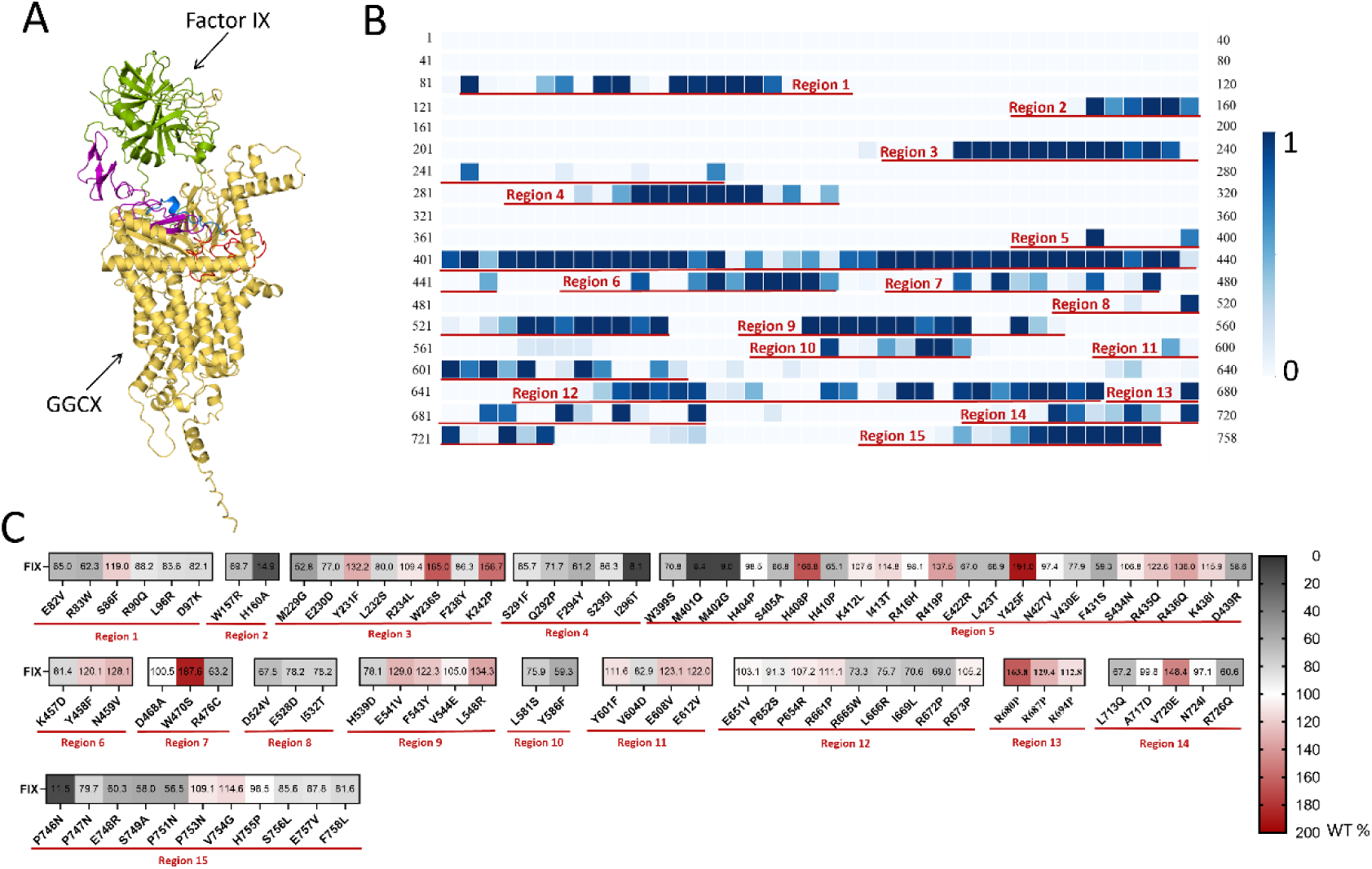
Prediction and validation the FIX-GGCX binding model. (**A**) AlphaFold3-predicted FIX-GGCX binding model. (**B**) Contact probability distribution of GGCX residues with FIX based on molecular dynamics simulation trajectory. The cut-off of 0.6 nm was used to access the contact between two residues. (**C**) Selection of specific points in each region of GGCX for validation (carboxylation activity). GGCX mutants were overexpressed into a GGCX-knockout cell line stably expressing FIX. After 48 hours of incubation in complete medium with 10 μM vitamin K1, carboxylation levels were quantified using ELISA.

Molecular dynamics simulation was used to assess the contact probability between GGCX residues and FIX. A heatmap (**Figure 1B**) was generated based on the final 60 ns simulation trajectory, indicating contact probabilities from 0 (no contact) to 1 (100% contact). Fifteen binding hotspots with significant interaction probabilities were identified, marked with red lines, and mapped onto the GGCX structure for further visualization (**Figures S1A-C**).

To experimentally validate the binding model, we performed mutagenesis on selected residues within the identified binding regions and evaluated their effects on GGCX carboxylation activity using cell-based assays.^23^ The carboxylation activity of each mutated residue was summarized in Figure 1C, highlighting those mutations in key binding regions significantly altered carboxylation efficiency. These findings confirm that multiple binding sites contribute to the overall interaction between GGCX and FIX.

GGCX binds to VKDPs via propeptides, anchoring the protein substrates to the enzyme and facilitating carboxylation^16^. We conducted a comprehensive analysis of GGCX’s interactions with FIX, identifying key binding sites in the GGCX-FIX propeptide interaction region using MM/PBSA binding energy calculations (**Figure 2A**). Critical residues, including Y231, H410, K412, N427, Y458, E528, F543, Y586, and E608, were found to be essential for these interactions. Binding strengths were visualized as a gradient from yellow (strong) to teal (weak). Subsequently, we examined the effects of mutations in these critical residues on FIX binding and carboxylation. Results from BiFC assay^14^ indicate that nearly all mutations decrease the binding affinity between GGCX and FIX (**Figure 2B**). However, most mutations, except for H410, Y586, and E528, slightly increased carboxylation activity without affecting reporter protein secretion (**Figure 2C and Figure S2**).

**Figure 2.**
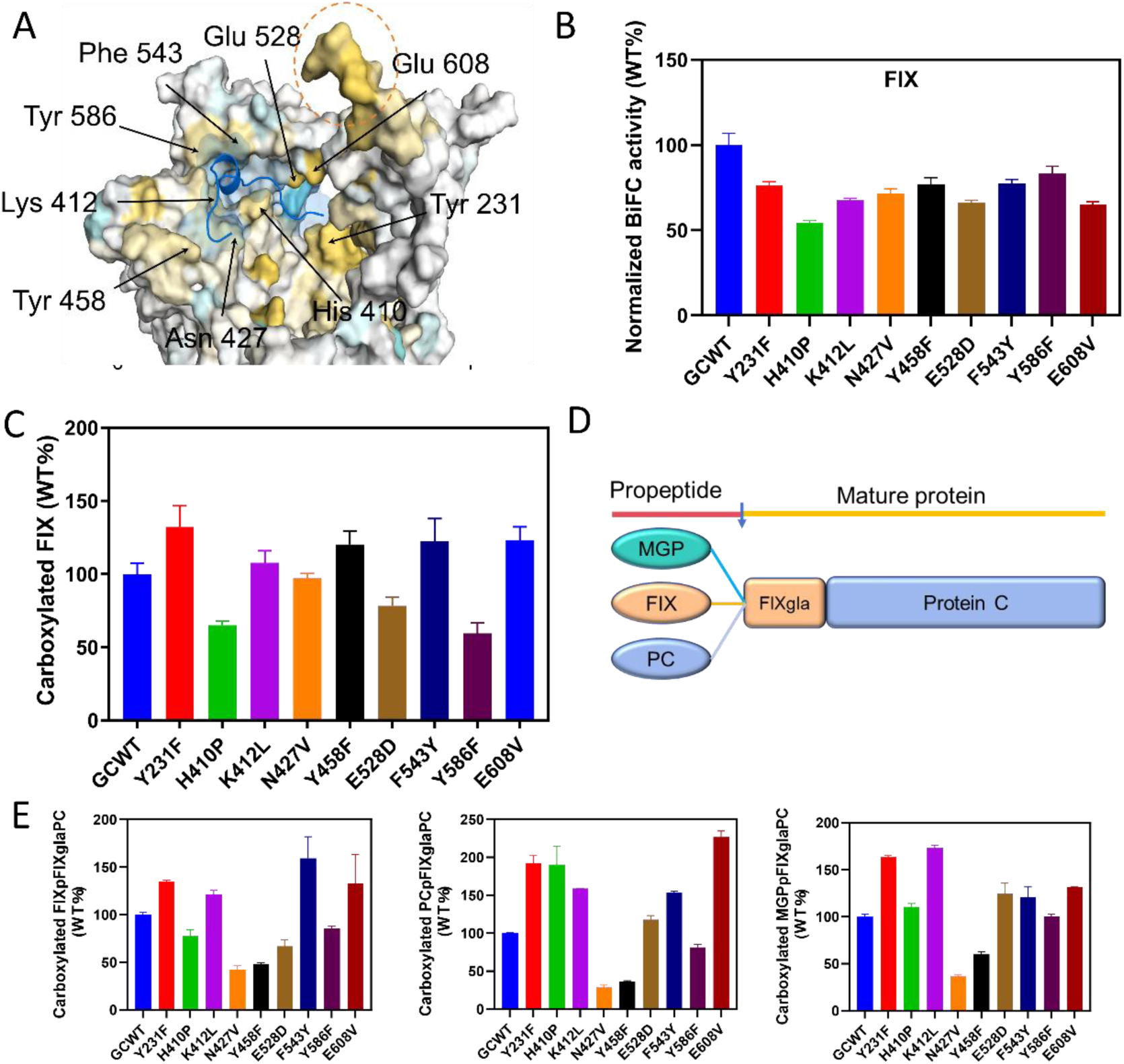
Binding of the FIX propeptide to GGCX depending on multiple amino acid sites. (**A**) Molecular visualization of binding sites between GGCX and FIX propeptide derived from MM/PBSA analysis. Teal indicates weaker binding, while yellow highlight stronger binding interactions. The red dashed circle highlights the C-terminal region of GGCX. (**B**) The interaction between FIX propeptide mutants and GGCX was examined in live cells without endogenous GGCX using the Venus-based BiFC assay. (**C**) Carboxylation activity of FIX under GGCX mutants. GGCX mutations were transfected into a GGCX-knockout cell line stably expressing the reporter-protein FIX. After 48 hours of incubation in complete medium with 10 μM vitamin K1, carboxylation levels were quantified using ELISA. (**D**) Topology diagram of reporter proteins guided by different propeptides. (**E**) Carboxylation activity of the three reporter proteins guided by FIX, MGP, and Protein C propeptides.

To further elucidate the role of the propeptide in GGCX binding, we designed three reporter proteins to mimic the binding behavior of the propeptide of FIX, MGP, and Protein C (**Figure 2D**). These reporter proteins exhibited varying affinities^16^ for GGCX (**Figure S3A**). The MGP propeptide-guided reporter showed the highest affinity, consistent with previous observations,^16^ but displayed lower carboxylation activity than the FIX-guided reporter (**Figure S3B and S3C**). The Protein C propeptide-guided reporter displayed intermediate binding affinity and reduced carboxylation activity (**Figure S3C**), supporting the hypothesis that distinct propeptides confer unique interaction profiles and enzymatic outcomes.

Next, we examined the effects of GGCX mutations on the carboxylation activity and binding affinity of these chimeric reporter proteins. Most mutations reduced the binding affinity of all three reporter proteins, except for Y586F (**Figure S3D**). Mutations at Y231, K412, F543, and E608 enhanced carboxylation across all reporters (**Figure 2E**), which correlated with their effects on FIX (**Figure 2B**). H410P and E528D mutations reduced carboxylation activity for FIX-directed reporters but enhanced or maintained activity for PC and MGP-directed reporters (Figure 2E). N427V and Y458F mutations decreased FIX carboxylation but increased activity for FIXglaPC reporters. Notably, the substrate-binding changes caused by GGCX mutations do not directly correlate with their carboxylation efficiency, possibly due to the balance between substrate binding and product release dynamics.^34,35^ Together, our findings demonstrate that multiple regions of GGCX converge to form a binding interface with critical residues essential for interacting with the propeptide.

### Variations in Vitamin K dosing for bleeding disorders stem from mutations in GGCX or the FIX propeptide region

Vitamin K is the mainstay treatment for VKCFD due to GGCX mutations^36^. Most GGCX mutations impair coagulation factor carboxylation at physiological vitamin K levels but show improved γ-carboxylation with high vitamin K1 (10 µM).^19^ Mutations in FIX propeptide can disrupt the interaction^20^ between GGCX and FIX, impairing FIX carboxylation and leading to bleeding disorders. However, high-dose vitamin K treatment is less commonly used for coagulation issues caused by FIX propeptide mutations, despite the similar pathogenic mechanism. To better understand how mutations in either GGCX or the FIX propeptide respond to vitamin K treatment, we introduced three mutations—A-10T and L-6R (pathogenic mutations), and F-16Y (a missense mutation)— at conserved sites in the FIX propeptide (**Figure S3A**)^20,37^. Our results showed that the A-10T, L-6R, and F-16Y mutations significantly reduced both coagulation and carboxylation activities of FIX due to impaired substrate binding without impact on their secretion (**Figure 3A-C and Figure S4A**).

**Figure 3.**
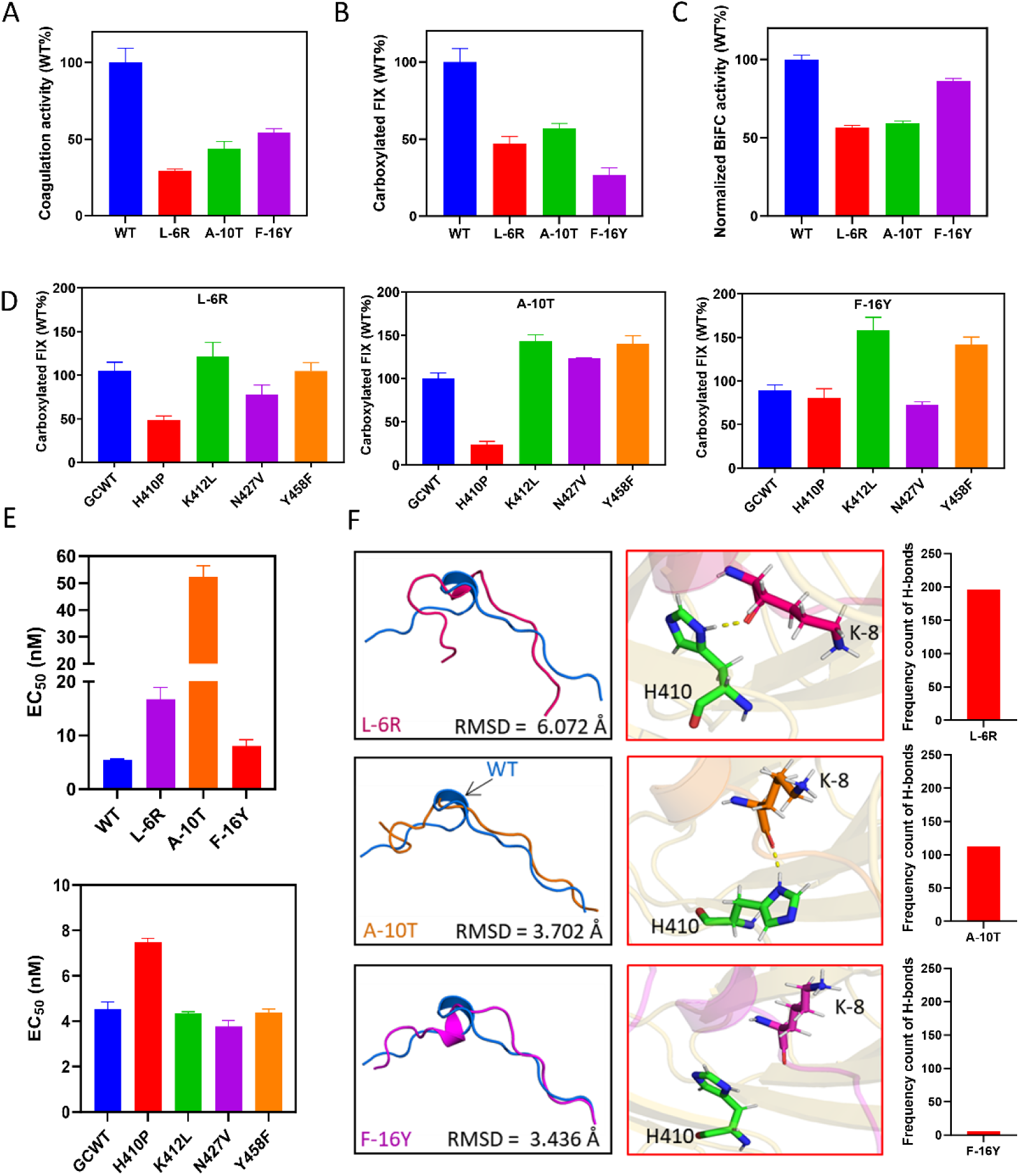
Vitamin K administration variations in bleeding disorders stem from mutations in GGCX or the FIX propeptide region. (**A**) Coagulation activity of FIX mutants was assessed using FIX-deficient plasma (SIEMENS). (**B**) Carboxylation activity of the three mutations was assessed by co-transfecting GGCX and the mutants into a GGCX-knockout cell line. After 48 hours of incubation in complete medium with 10 μM vitamin K1, carboxylation levels were quantified using ELISA. (**C**) The interaction between FIX propeptide mutants and GGCX was examined in live cells without endogenous GGCX using the Venus-based BiFC assay. (**D**) Effect of FIX’s propeptide mutations on carboxylation activity under GGCX mutations. The FIX mutations, along with GGCX and its mutants, were co-transfected into a GGCX-knockout cell line. After 48 hours in complete medium with 10 μM vitamin K1, carboxylation activity was measured by ELISA. (**E**) Vitamin K concentration titration for FIX carboxylation under various GGCX mutations, as well as the EC_50_ values of FIX propeptide mutants under both wild-type and mutated GGCX. GGCX or mutants were transiently co-expressed with various reporter-protein in GGCX-knockout cell line. Then the reporter proteins’ carboxylation was accessed as described in Figure 1F. (**F**) Visualization of altered binding patterns by the propeptide mutants. This panel shows how mutations in the FIX propeptide alter its conformation (black boxes) and binding patterns (red boxes) with GGCX, affecting enzyme interactions. The histogram represents the number of hydrogen bonds that occur in 500 frames of MD simulations.

Given the differential carboxylation effects of H410P, K412L, N427V, and Y458F mutations on various reporter proteins (**Figure 2E**), we selected these mutations to assess their impact on FIX propeptide mutants. GGCX mutants had no significant effect on reporter proteins secretion (**Figure S4B-D**). The H410P mutation significantly impaired carboxylation efficiency in both the L-6R and A-10T mutants, while the K412L mutation consistently enhanced carboxylation across all mutant variants (**Figure 3D**). The Y458F mutation led to a moderate increase in carboxylation in the A-10T and F-16Y mutants (**Figure 3D**). We further measured the EC_50_ values of vitamin K1 for FIX carboxylation under various GGCX mutations, as well as the EC_50_ values of FIX propeptide mutants under both wild-type and mutated GGCX. Figure 3E illustrates that propeptide mutations, such as A-10T, result in a significant 10-fold increase in the EC_50_ of vitamin K, consistent with previous findings.^34^ In contrast, GGCX mutations, like H410P, cause a modest, approximately two-fold increase in EC_50_ (**Figure 3E**). These results align with prior studies^14^, which suggest that most pathogenic GGCX mutations lead to only minor or moderate variations in the EC_50_ of vitamin K.

We hypothesize that the difference arises from the propeptide’s short length (18 amino acids) and inherent flexibility, where a single mutation induces a substantial conformational change, disrupting substrate recognition and binding, as supported by molecular dynamics simulations (**Figure 3F**). In the binding simulations of A-10T and L-6R mutants, the H410 residue of GGCX forms stable hydrogen bonds with the K-8 residue of the propeptide (**Figure 3F**). However, this interaction is weaker in the F-16Y mutant, as indicated by the lower hydrogen bond frequency in Figure 3F. Hydrogen bonding disruptions affect GGCX and FIX propeptide interactions, correlating with observed carboxylation changes in these mutations. Figure S5 shows a similar trend, where the N427 residue of GGCX forms stable hydrogen bonds with the V-17 residue of the propeptide in the F-16Y and L-6R mutants. Disruptions at these sites weaken hydrogen bonds and affect GGCX-FIX binding. Binding strength changes correspond to carboxylation alterations from propeptide mutations. Additionally, Figures S5 indicate that binding strength variations for propeptide mutants are mainly affected by changes in hydrophilic/hydrophobic and π–π interactions between the conserved residues (N-9, V-17, and F-16) in propeptide and the paired residues in GGCX (K412, N427, and Y458).

Collectively, mutations in the FIX propeptide more severely affect GGCX-FIX binding than GGCX mutations. High-dose vitamin K therapy partially restores carboxylation in GGCX mutants, but is less effective in propeptide mutants.

### GGCX’s C terminal is vital for VKDPs’ differentially binding and carboxylation

The C-terminal region of GGCX plays a critical role in substrate affinity and recognition (**Figure 1C**).^14^ Using molecular dynamics simulations, we explored the functional significance of GGCX’s C-terminal region and identified key interactions with FIX’s functional domain (**Figure 4A**). These simulations highlighted critical residues and structural features in the C-terminal region essential for GGCX’s interaction with FIX. To assess the impact of GGCX C-terminal mutations on VKDP carboxylation, we utilized three reporter proteins—FIX, FIXglaPC, and Protein C (**Figure 4B**)—which share structural similarities but possess distinct functional domains. Mutations in the H755-F758 region of GGCX led to an approximate 30% reduction in the carboxylation activity of Protein C and FIXglaPC, with the E757V mutation causing a more substantial 50% decrease in Protein C carboxylation (**Figure 4C**). In contrast, FIX carboxylation was largely unaffected by these mutations. Reporter protein expression levels remained consistent, except for Protein C (**Figure S6A**), which exhibited reduced secretion due to undercarboxylation.^38^ The primary structural difference between FIX and FIXglaPC is their functional domains: FIXglaPC shares its functional domain with Protein C, while FIX possesses a unique domain (**Figure 4B**). These structural variations likely influence binding affinity to GGCX, thereby impacting carboxylation efficiency.

**Figure 4.**
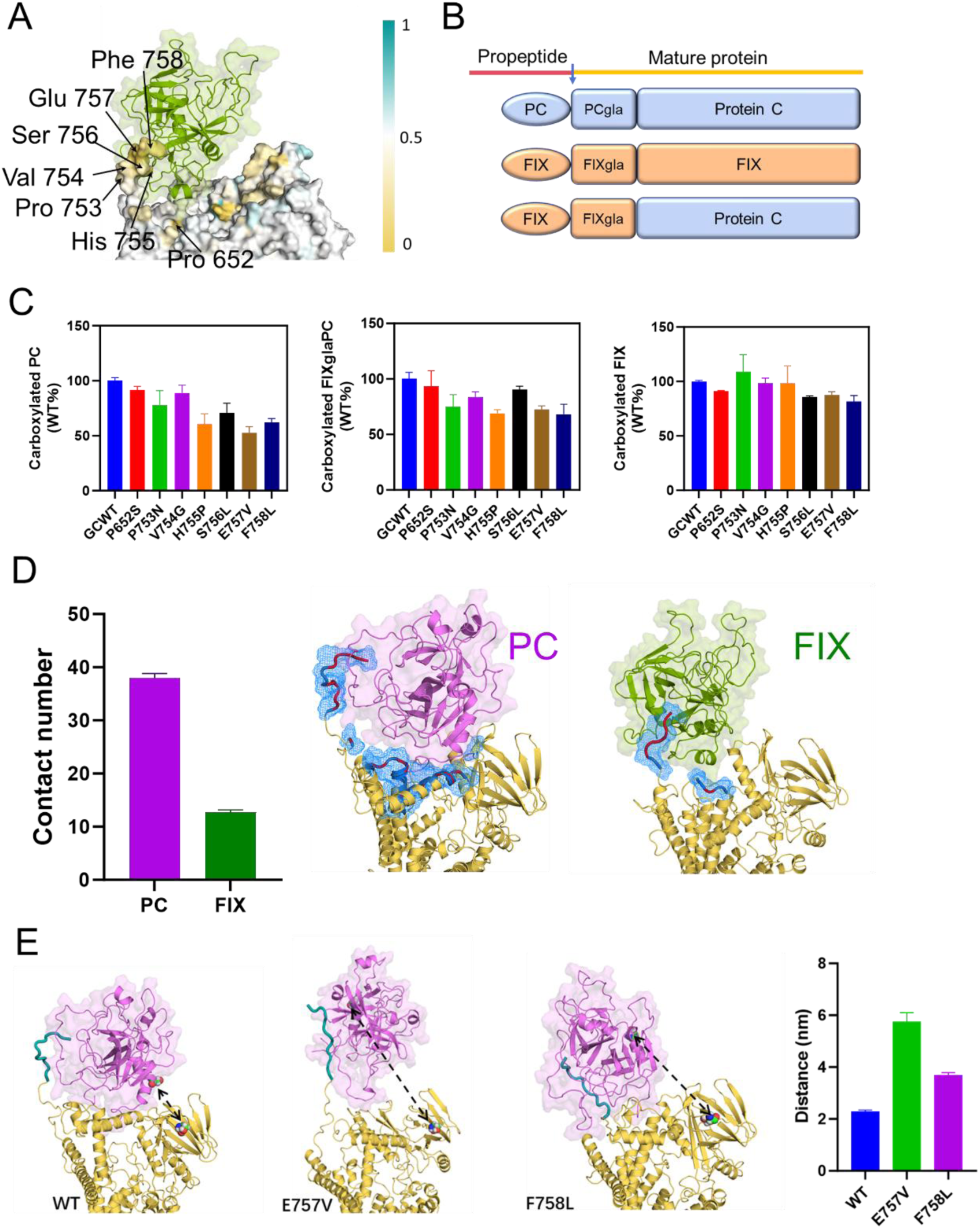
GGCX’s C-terminal is vital for VKDPs’ differentially binding and carboxylation. (**A**) Molecular visualization of binding sites between GGCX and FIX’s functional domain. The color scale shows binding intensity: teal indicates weak interactions, and yellow indicates strong ones. (**B**) Topology diagram of distinct reporter proteins. (**C**) Carboxylation activity of reporter proteins with GGCX mutations, described in Figure 1C. (**D**) Contact number of GGCX residues and molecular visualization of GGCX binding with functional domains of Protein C and FIX. Light blue highlights potential interaction regions between GGCX and FIX’s functional domain, while light red indicates key binding sites with Protein C or FIX’s functional domains. (**E**) The structural modeling and quantitative analysis demonstrate how the E757V and F758L mutations in GGCX’s C-terminal region alter the spatial arrangement of the Protein C functional domain. The distance between two selected residues in GGCX (Ser552) and PC (Asp401) was calculated based on simulation trajectories.

We then experimentally assessed reporter proteins binding affinity to GGCX using a BiFC assay. Binding affinity for the FIX reporter remained largely unchanged, while Protein C and FIXglaPC showed slightly reduced binding (less than 20%), particularly in mutants E757V and F758L (**Figure S6B**). Notably, the reduction in binding strength for Protein C and FIXglaPC was smaller than the corresponding decrease in carboxylation activity (**Figure 4C and S6B**). The P652S mutation increased binding affinity with Protein C and FIXglaPC but did not significantly enhance their carboxylation activity. These results suggest that although VKDP functional domains interact with GGCX’s C-terminal region, changes in binding affinity do not always result in proportional changes in carboxylation efficiency.

We further analyzed the binding sites between GGCX and the functional domains of FIX and Protein C using molecular dynamics simulations (**Figure 4D**). Both statistical and visualized results showed stronger interactions between the C-terminal region of GGCX and the functional domain of Protein C compared to FIX. Then, we examined the binding conformations of Protein C’s functional domain with GGCX, focusing on mutations at H755, E757, and F758. Virtualizational structure (**Figure 4E**) revealed significant conformational changes in these mutants compared to wild-type GGCX. This aligns with previous findings indicating substrate-induced reorientation of the C-terminal region of GGCX.^39^ To quantify these differences, the distances between Asp401 in Protein C’s functional domain and Ser552 in GGCX were calculated. Mutations at H755, E757, and F758 increased the distances (**Figure 4E and S7**), potentially triggering a cascade effect that disturbs residue binding in other domains. These effects may result from either the close spatial proximity of these regions within VKDP or an allosteric mechanism connecting them^35,40–42^, particularly involving the Gla domain. These findings reinforce that GGCX’s C-terminal region differentially impacts the binding and carboxylation of VKDPs, which possess unique functional domains.

### Localization and characterization of the active site of Vitamin K epoxidation in GGCX

Understanding GGCX’s active sites is vital for clarifying the molecular mechanisms underlying severe bleeding disorders. This study investigates GGCX’s epoxidase activity, crucial for VKDP carboxylation. We studied the interaction between GGCX and reduced vitamin K (KH₂) through molecular docking and molecular dynamics simulations. Theoretical analysis identified five key binding regions in GGCX essential for KH₂ binding (**Figure 5A and Figure S8**). Molecular visualization further elucidated the GGCX-KH₂ binding mode (**Figure 5B**). The effects of critical GGCX residues on enzymatic activity were assessed using three reporter proteins: FIX, FIXglaPC, and Protein C. Mutations at these residues abolished carboxylation activity across all reporter proteins, except for the Y211 mutation, which had a milder impact (around 50% reduction for FIX’s carboxylation) (**Figure 5C**). These mutations did not affect GGCX expression levels, except for K218 and M303, which showed moderate reduced expression^43^ (**Figure 5D**), suggesting that these residues primarily affect enzymatic function or substrate affinity rather than protein stability. Using BiFC analysis, we found that the afore mentioned GGCX mutations had no significant effect on substrate binding, except for K218A and M303R (**Figure S9**). The reduced binding observed for these two mutants is consistent with the decreased protein expression in Figure 5D.

**Figure 5.**
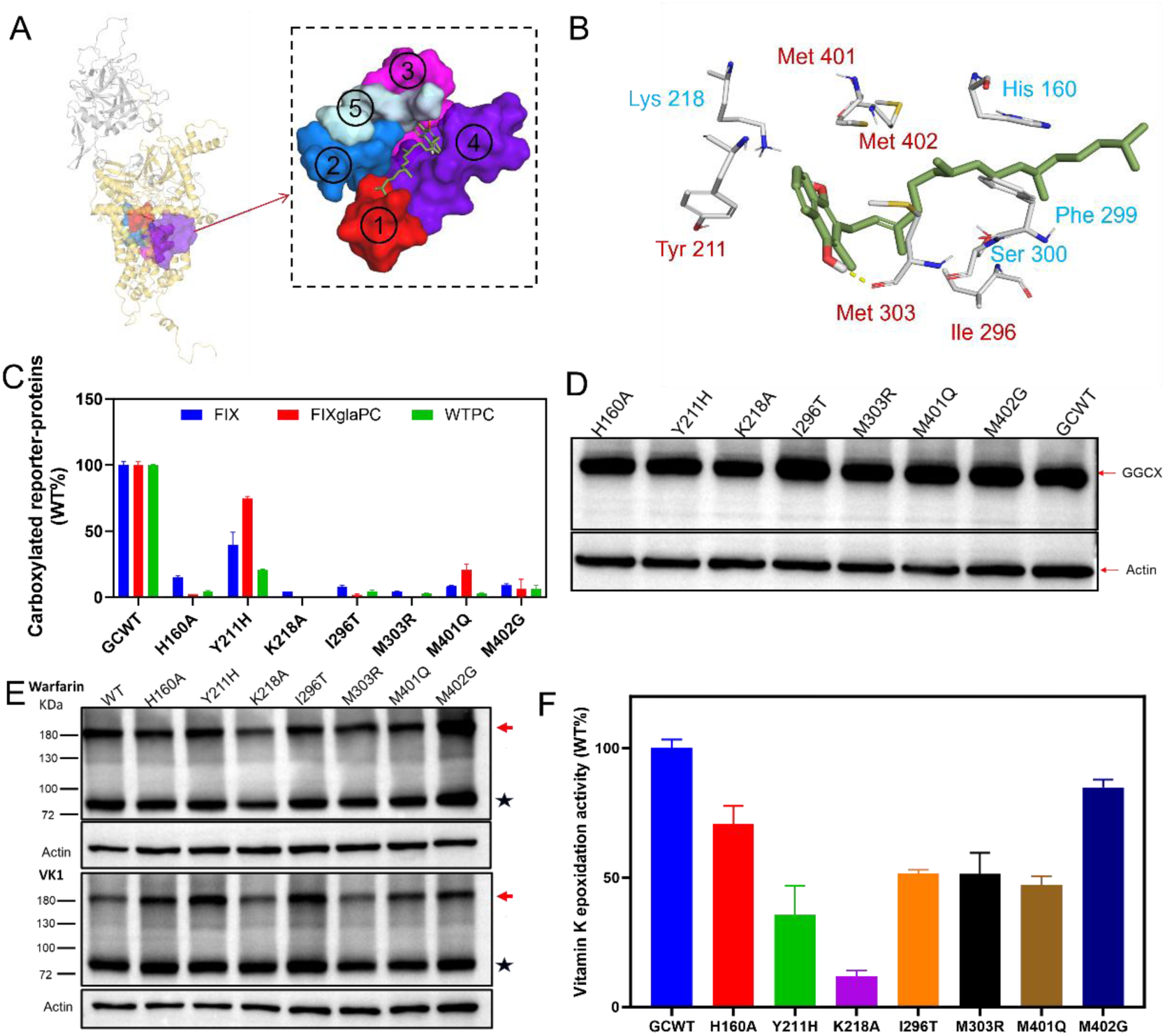
Localization and characterization of the active site of Vitamin K epoxidation in GGCX. (**A**) The visualization shows five distinct strong binding regions of GGCX with reduced vitamin K1. (**B**) Key residues in GGCX-KH_2_ binding. Reported residues were colored in blue. (**C**) The carboxylation activity of FIX under different GGCX mutations. GGCX/specific mutations were transfected into a GGCX-knockout cell line stably expressing the reporter protein FIX. After 48 hours in complete medium with 10 μM vitamin K1, carboxylation levels were quantified by ELISA. (**D**) Expression of GGCX mutants with western blot. GGCX and mutants were transiently expressed in HEK293 without endogenous GGCX cell line. After 48 hours incubation, cell lysate was analyzed with western blot using a rabbit anti-GGCX polyclonal antibody. (**E**) The interaction between GGCX and its protein substrate examined in live cells using DSS (disuccinimidyl suberate) cross-linking. Wild-type GGCX and mutations were co-expressed with FIXgla-MBP in GGCX-deficient HEK293 cells. Forty-eight hours later, transfected cells were crosslinked with 4 mM DSS for 30 minutes. The whole-cell lysate was analyzed by Western blot using rabbit anti-GGCX polyclonal antibody. The GGCX and FIXgla-MBP cross-linked band was marked with a red arrow. The GGCX is marked with a star. (**F**)The effects of GGCX mutations on vitamin K epoxidation activity. Wild-type GGCX and its mutant were transiently expressed in GGCX-deficient HEK293 cells. Transfected cells were incubated with vitamin K, and the conversion to vitamin K epoxide by GGCX was measured using reversed-phase high-performance liquid chromatography.

Furthermore, we employed a combination of DSS (disuccinimidyl suberate) chemical cross-linking^14^ and a cell-based vitamin K epoxidase activity assay^33^ to systematically evaluate the effects of the aforementioned mutations on substrate binding and GGCX enzymatic activity. We co-expressed GGCX/mutants with reporter proteins in GGCX-deficient HEK293 cells to study enzyme-substrate interactions. DSS cross-linking studies in GGCX-deficient HEK293 cells showed high cross-linking efficiency between GGCX mutants and VKDPs, similar to wild-type GGCX with warfarin. Most GGCX mutants can partially dissociate VKDPs with vitamin K, but less efficiently than the wild type (**Figure 5E**). This observation supports the carboxylation assay results (**Figure 5C**), showing that these mutations lead to a loss of GGCX activity.

We further evaluated the impact of GGCX mutations on the vitamin K epoxidation activity. The K218A mutation nearly eliminated epoxidase activity (**Figure 5F**), while the H160A, I296T, M303R, and M401Q mutations retained approximately 50% of epoxidase activity but showed no carboxylation activity (**Figures 5C and 5F**). The Y211H mutation reduced epoxidase activity by 60% and carboxylation activity by 50%, indicating a primary effect on epoxidase function. The M401Q mutation led to a 16% decrease in epoxidase activity, suggesting a notable impact on carboxylation. Collectively, the residues H160, Y211, I296, K218, M303, M401, and M402 are essential for vitamin K epoxidation, while H160, I296, M303, M401, and M402 are also critical for carboxylation.

### The carboxylation and vitamin K epoxidation active centers partially overlap

Previous study has demonstrated that BGP interact with GGCX through its Gla domain^26^. To identify potential binding sites on GGCX involved in VKDP interaction and carboxylation, we analyzed simulation results and highlighted residues on GGCX that may interact with the Gla domains of FIX and BGP (**Figure 6A, Figure S10**). These findings suggest that GGCX utilizes similar binding sites to interact with the Gla domains of BGP and FIX, including residues R90, M229, R234, Q292, F294, I296, F299, S300, R435, R436, R665, and R672. Notably, F299, S300^14^, and I296 (**Figure 5F**) have been confirmed as essential for vitamin K epoxidation activity. Given their spatial proximity, we hypothesize that the active sites for GGCX’s carboxylation and vitamin K epoxidation activities may partially overlap.

**Figure 6.**
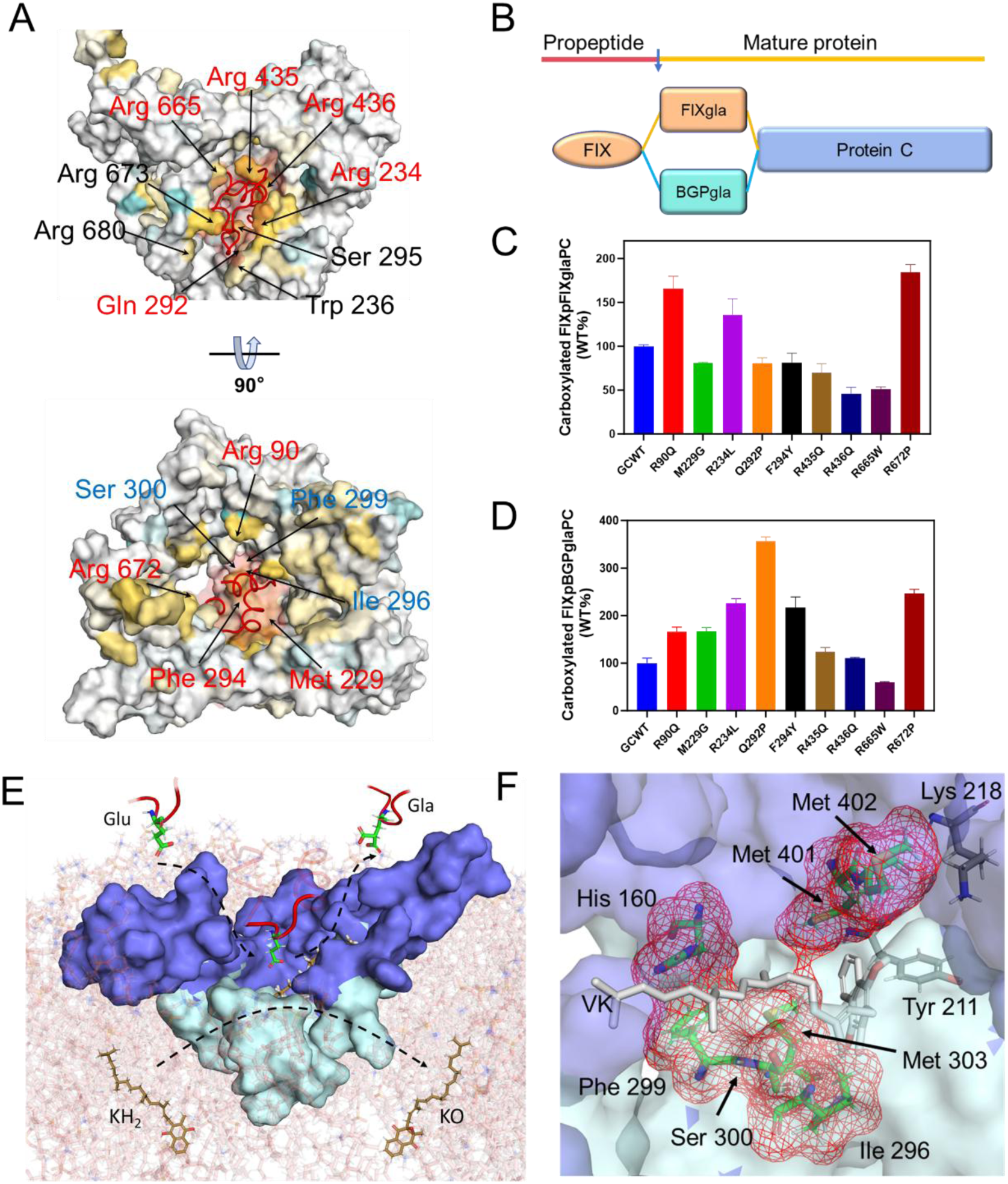
Carboxylation and vitamin K epoxidation active centers partially overlap. (**A**) Molecular visualization of the key binding residues of GGCX with the Gla domain of FIX. Residues colored blue are related to GGCX’s enzymatic activity. Residues highlighted in red represent conserved binding sites that interact with the Gla domains of both BGP and FIX. (**B**) Topology diagram of reporter proteins with different Gla domains, shows two reporter proteins with distinct gla domains that share the same propeptide and functional domains. (**C**) and (**D**) depict the carboxylation activity of the two reporter proteins under various GGCX mutations. GGCX and specific mutations with reporter proteins were co-expressed in a GGCX-knockout cell line. After 48 hours in complete medium with 10 μM vitamin K1, carboxylation levels were measured by ELISA. (**E**) Molecular visualization illustrates the coupling between VKDP carboxylation and the vitamin K epoxidation. (**F**) Molecular visualization displays partially overlapping active sites of carboxylation and vitamin K epoxidation. The Gla binding regions are colored in slate. And the residues related to vitamin K epoxidation are colored in palecyan.

To determine the role of these residues in carboxylation, individual mutations were analyzed using reporter proteins with identical propeptide and functional domains but distinct Gla domains (**Figure 6B**). Figures 6C and 6D show that mutations at R90, R234, and R672 enhance carboxylation activity in both reporter proteins, while the R665 mutation reduces activity. Residues M229, Q292, and F294 had minimal impact on the FIX Gla domain reporter but significantly increased carboxylation of the BGP Gla domain. Similarly, mutations at R435 and R436 decreased FIX Gla domain carboxylation without affecting BGP carboxylation. Notably, these mutations had no significant impact on the binding of BGP (**Figure S11**). Glu-containing peptides likely play a critical role in carboxylation by stabilizing the propeptide-carboxylase complex.^42^ We hypothesize that these GGCX mutations specifically disrupt local interactions with key residues of VKDPs’ Gla domain, thereby affecting carboxylation. Molecular analysis further identified distinct VKDP binding residues associated with these GGCX sites, highlighting their differential binding (**Figure S12**).

As shown in visualized structure of simulation conformation, the regions involved in binding the VKDP Gla domain form a circular cavity essential for carboxylation (**Figure 6A and Figure S13**). This cavity is spatially connected to the vitamin K epoxidation center (**Figure 6E and S13**). Overlapping residues (H160, I296, F299, S300, M303, M401, and M402) participate in both enzymatic functions, suggesting that the carboxylation active center is adjacent to the epoxidation center (**Figure 6E**). Building on this, we visualized the molecular structure illustrating the coupling between VKDP carboxylation and vitamin K epoxidation. Figure 6E suggests that VKDPs, directed by their propeptides, bind to GGCX with their Gla domains positioned within a binding cavity. The adjacent vitamin K epoxidation center oxidizes reduced vitamin K (KH₂) to its epoxidized form (KO), supplying the energy required for the processive^42^ carboxylation of glutamate residues (Glu) on VKDPs.

## Discussion

The γ-carboxylation of VKDPs is a critical post-translational modification catalyzed by GGCX. Disruption of this process results in dysfunctional VKDPs, leading to bleeding disorders, osteoporosis, and vascular calcification. Despite its importance, the molecular mechanisms underlying GGCX’s substrate binding and carboxylation remain poorly understood due to limited structural data. This study combines computational and experimental approaches to elucidate the binding model that integrates GGCX, FIX, and reduced vitamin K (VK) (**Figure 7A**), providing new insights into substrate recognition and enzymatic function. Visualization structure shows the spatial orientation of FIX interacting with GGCX, while the reduced vitamin K molecule positioning near the enzyme’s active site (**Figure 7 A and 6E**).

**Figure 7.**
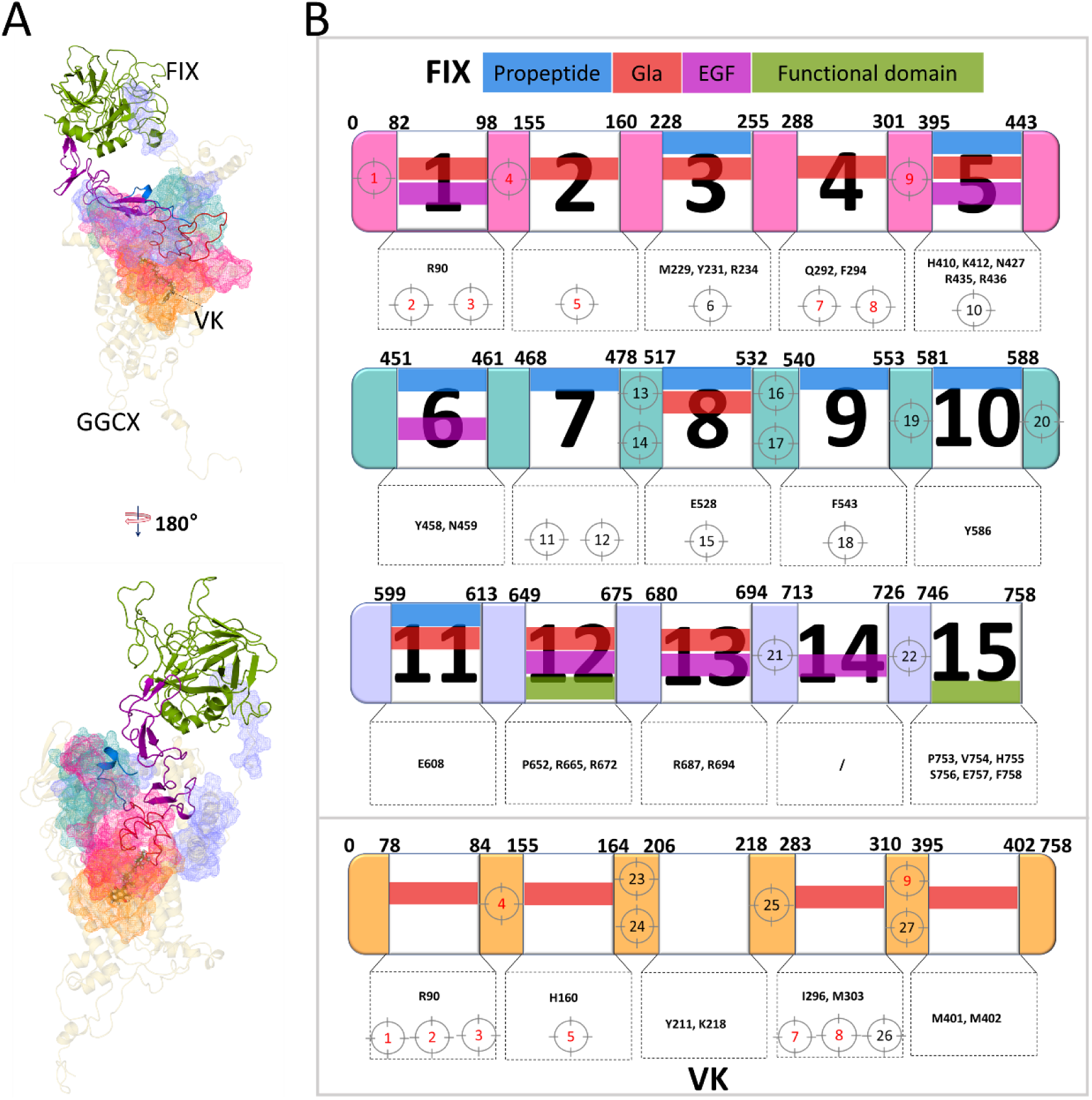
Validation of the FIX-GGCX binding and carboxylation model in the bleeding disorders. (**A**) Visualized structure of the FIX-GGCX binding model. (**B**) Validation of the GGCX-FIX binding and carboxylation model using GGCX mutations associated with VKCFD1 disease. Pathogenic mutations are numbered targets, highlighted in red for those conserved in their interaction with Factor IX (FIX) and reduced vitamin K. A detailed description of these mutations can be found in Supplementary Table 1. Additionally, pathogenic mutations located near the binding regions, within 10 residues, are also included in this analysis. The residues highlighted in black within the dashed box represent sites that have been experimentally validated as involved in substrate recognition, binding, or catalytic activity.

Our data reveal that multiple sites and regions on GGCX synergistically contribute to its recognition of VKDP propeptides and its binding to VKDPs (**Figure 1B and Figure 2A**). Clinical evidence supports this conclusion (**Figure 7B**), as GGCX pathogenic mutations in these regions, such as V255M^27^, H404P^44^, L394R^17^, R476C/H^24^, I532T^45^, and T591K^46^ have been associated with reduced carboxylation activity due to impaired substrate binding affinity. These pathogenic mutations are dispersed across multiple regions of GGCX, and the currently identified propeptide-binding sequence (AA495-513)^47^ does not sufficiently explain this phenomenon. Our results indicate that multiple regions of GGCX converge to form a binding interface containing critical residues essential for interacting with the FIX propeptide (**Figure 2A**). Dispersed pathogenic mutants may further disrupt the structural integrity of this interface, compromising substrate recognition and binding. Previous experimental results further support this observation, showing that multiple regions of GGCX exhibit indirect cooperative structural dependency in substrate binding^39^, when they do not directly contribute to propeptide recognition. Additionally, prior study indicates that substrates binding to the active site modify the propeptide binding site of carboxylase.^41^ These observations validates our proposed model, providing a mechanistic explanation for the dispersed distribution of pathogenic mutations across various regions of GGCX and their collective impact on substrate binding.

The identification of the vitamin K protein-interacting sequence (VKS)^48^ and glutamate-binding sites^49^ underscores the critical role of non-propeptide binding regions in GGCX-substrate interaction. Consistent with these findings, our results reveal that mutations in GGCX’s C-terminal regions (**Figure 4**) and in residues involved in Gla domain binding (**Figure 6**) have minimal impact on the overall binding affinity between GGCX and VKDPs (**Figure S6B and S11**). However, these mutations do affect local interactions with specific VKDP residues (**Figure 4E and S12**), affecting VKDP’s carboxylation. These non-propeptide regions likely play a crucial role in stabilizing the GGCX-VKDP complex during the enzymatic process, thereby ensuring processive carboxylation. This aligns with previous studies that demonstrated significant binding energy between the Glu residues of VKDPs and the active site of GGCX ^50,51^.

Our findings indicate that the structural variations within VKDPs modulate their binding affinity and carboxylation efficiency (**Figure 4**). This differential substrate recognition mechanism may explain the diverse clinical manifestations of GGCX mutations, including coagulation disorders, non-coagulation symptoms, or a combination of both.^4,24,27^ Meanwhile, the EGF domain^52^ of FIX connects the Gla domain to the functional domain and interacts strongly with GGCX residues R90 and R435, which also bind to the Gla domain (**Figure 6A and Figure S14**). Additionally, residues N427 and Y458 of GGCX interact with the EGF domain while simultaneously participating in propeptide binding (**Figure 2A**), further underscoring the synergistic contribution of multiple sites and regions to the GGCX-FIX interaction.

Our work identified five novel active-site residues—Y211, I296, M303, M401, and M402—essential for GGCX’s dual enzymatic functions (**Figure 5**), in addition to previously characterized residues (F299, S300, H160, and K218).^34,43,53^ These residues participate in both carboxylation and vitamin K epoxidation, refining our understanding of GGCX’s active center. And the M401 and M402 residues are located in the glutamate-binding region.^49^ While these two residues appear critical for catalytic activity (**Figure 5C and 5F**), BiFC assays (**Figure S9**) indicate that mutations at M401 and M402 do not significantly impact VKDP binding, suggesting a primary role in enzymatic catalysis rather than substrate interaction.

Based on the spatial connectivity between the carboxylation and vitamin K epoxidation active centers (**Figure 5B and 6A**), we propose a model for GGCX-mediated carboxylation of VKDPs (**Figure 6E**). This model ensures efficient coupling of carboxylation and vitamin K epoxidation^53^, aligning with the “base strength amplification mechanism” proposed by Dowd et al.^54^, where the oxygenation of reduced vitamin K generates free energy to transform a weak base into a strong base, facilitating glutamate deprotonation. Meanwhile, Rishavy et al. proposed that K218 deprotonates KH₂ to start carboxylation, with H160 crucial for carbanion formation.^43,53^ These two residues are central to our carboxylation model (**Figure 6E**), reinforcing its validity. Moreover, the active cavity formed by these critical residues is partially embedded within the third transmembrane region of GGCX, oriented toward the ER lumen^55^ (**Figure 6E**). This localization is consistent with the hydrophobic nature of vitamin K and supports the understanding that carboxylation occurs within the ER lumen.^56^

In summary, our study emphasizes the highly regulated and dynamic nature of the γ-carboxylation process, which relies on precise coordination among multiple regions of GGCX. The proposed GGCX-FIX binding and carboxylation model aligns with known pathogenic mutations in GGCX, highlighting its potential for elucidating carboxylation defects and advancing the understanding of coagulation disorders.

## Acknowledgments

This work are supported by grants 82370136 (to Z.H.) from National Natural Science Foundation of China; grant HL131690 (to J.K.T. and D.W.S.) from the National Institutes of Health, USA; grant BK20231333 (to Z.H.) and BK20210828 (to S.L.) from Natural Science Foundation of Jiangsu Province; Jiangsu Specially-Appointed Professor Start-up Funds grant (to Z.H.); and grant 2023M732983 (to S.L.), 2022M712691 (to N.J.) from China Postdoctoral Science Foundation.

## Authorship Contributions

Z. H., J. T., and J. H. conceived the studies. K. L. and M. H. performed the cell-based functional studies, K. L., L. C. and D. H. performed the BiFC assay, S. L., J. T. and Y. J performed MD simulation and Molecular docking, K. L., D. L. constructed plasmids and created GGCX mutations., Y. G. performed the chemical cross-linking, K. L. and Y. Z. performed epoxidase activity assays. Z. H., J. T., J. H., and S. L. wrote the manuscript, analyzed the data and wrote the manuscript. All authors reviewed and contributed to the manuscript.

## References

1. Tie JK, Stafford DW. Structural and functional insights into enzymes of the vitamin K cycle. J Thromb Haemost. 2016;14(2):236–247.

2. Berkner KL, Runge KW. Vitamin K-Dependent Protein Activation: Normal Gamma-Glutamyl Carboxylation and Disruption in Disease. Int J Mol Sci. 2022;23(10).

3. Girolami A, Ferrari S, Cosi E, Santarossa C, Randi ML. Vitamin K-Dependent Coagulation Factors That May be Responsible for Both Bleeding and Thrombosis (FII, FVII, and FIX). Clin Appl Thromb Hemost. 2018;24(9_suppl):42S–47S.

4. Ghosh S, Oldenburg J, Czogalla-Nitsche KJ. The Role of GRP and MGP in the Development of Non-Hemorrhagic VKCFD1 Phenotypes. Int J Mol Sci. 2022;23(2):798.

5. Schurgers LJ, Uitto J, Reutelingsperger CP. Vitamin K-dependent carboxylation of matrix Gla-protein: a crucial switch to control ectopic mineralization. Trends Mol Med. 2013;19(4):217–226.

6. Mishima E, Wahida A, Seibt T, Conrad M. Diverse biological functions of vitamin K: from coagulation to ferroptosis. Nat Metab. 2023;5(6):924–932.

7. Li WK, Schulman S, Dutton RJ, Boyd D, Beckwith J, Rapoport TA. Structure of a bacterial homologue of vitamin K epoxide reductase. Nature. 2010;463(7280):507–U123.

8. Mladenka P, Macakova K, Kujovska Krcmova L, et al. Vitamin K – sources, physiological role, kinetics, deficiency, detection, therapeutic use, and toxicity. Nutr Rev. 2022;80(4):677–698.

9. Liu S, Li S, Shen G, Sukumar N, Krezel AM, Li W. Structural basis of antagonizing the vitamin K catalytic cycle for anticoagulation. Science. 2021;371(6524):eabc5667.

10. De Vilder EY, Debacker J, Vanakker OM. GGCX-Associated Phenotypes: An Overview in Search of Genotype-Phenotype Correlations. Int J Mol Sci. 2017;18(2):240.

11. Dalmeijer GW, van der Schouw YT, Magdeleyns EJ, et al. Circulating desphospho – uncarboxylated matrix γ-carboxyglutamate protein and the risk of coronary heart disease and stroke. J Thromb Haemost. 2014;12(7):1028–1034.

12. Watzka M, Geisen C, Scheer M, et al. Bleeding and non-bleeding phenotypes in patients with GGCX gene mutations. Thromb Res. 2014;134(4):856–865.

13. Ghosh S, Kraus K, Biswas A, et al. GGCX variants leading to biallelic deficiency to gamma-carboxylate GRP cause skin laxity in VKCFD1 patients. Hum Mutat. 2022;43(1):42–55.

14. Hao Z, Jin DY, Chen X, Schurgers LJ, Stafford DW, Tie JK. gamma-Glutamyl carboxylase mutations differentially affect the biological function of vitamin K-dependent proteins. Blood. 2021;137(4):533–543.

15. Ayombil F, Camire RM. Insights into vitamin K-dependent carboxylation: home field advantage. Haematologica. 2020;105(8):1996–1998.

16. Stanley TB, Jin DY, Lin PJ, Stafford DW. The propeptides of the vitamin K-dependent proteins possess different affinities for the vitamin K-dependent carboxylase. J Biol Chem. 1999;274(24):16940–16944.

17. Brenner B, Sanchez-Vega B, Wu SM, Lanir N, Stafford DW, Solera J. A missense mutation in gamma-glutamyl carboxylase gene causes combined deficiency of all vitamin K-dependent blood coagulation factors. Blood. 1998;92(12):4554–4559.

18. Mutucumarana VP, Stafford DW, Stanley TB, et al. Expression and Characterization of the Naturally Occurring Mutation L394R in Human γ-Glutamyl Carboxylase. J Biol Chem. 2000;275(42):32572–32577.

19. Ghosh S, Kraus K, Biswas A, et al. GGCX mutations show different responses to vitamin K thereby determining the severity of the hemorrhagic phenotype in VKCFD1 patients. J Thromb Haemost. 2021;19(6):1412–1424.

20. Oldenburg J, Quenzel EM, Harbrecht U, et al. Missense mutations at ALA-10 in the factor IX propeptide: an insignificant variant in normal life but a decisive cause of bleeding during oral anticoagulant therapy. Br J Haematol. 1997;98(1):240–244.

21. Belvini D, Salviato R, Radossi P, et al. Molecular genotyping of the Italian cohort of patients with hemophilia B. Haematologica. 2005;90(5):635–642.

22. Chu K, Wu SM, Stanley T, Stafford DW, High KA. A mutation in the propeptide of Factor IX leads to warfarin sensitivity by a novel mechanism. J Clin Invest. 1996;98(7):1619–1625.

23. Tie JK, Carneiro JDA, Jin DY, Martinhago CD, Vermeer C, Stafford DW. Characterization of vitamin K-dependent carboxylase mutations that cause bleeding and nonbleeding disorders. Blood. 2016;127(15):1847–1855.

24. Vanakker OM, Martin L, Gheduzzi D, et al. Pseudoxanthoma elasticum-like phenotype with cutis laxa and multiple coagulation factor deficiency represents a separate genetic entity. J Invest Dermatol. 2007;127(3):581–587.

25. Darghouth D, Hallgren KW, Issertial O, et al. Compound Heterozygosity of a W493C Substitution and R704/Premature Stop Codon within the gamma-Glutamyl Carboxylase in Combined Vitamin K-Dependent Coagulation Factor Deficiency in a French Family. Blood. 2009;114(22):533–534.

26. Houben RJ, Jin D, Stafford DW, et al. Osteocalcin binds tightly to the gamma-glutamylcarboxylase at a site distinct from that of the other known vitamin K-dependent proteins. Biochem J. 1999;341 (Pt 2):265–269.

27. Li QL, Grange DK, Armstrong NL, et al. Mutations in the GGCX and ABCC6 Genes in a Family with Pseudoxanthoma Elasticum-Like Phenotypes. J Invest Dermatol. 2009;129(3):553–563.

28. Abramson J, Adler J, Dunger J, et al. Accurate structure prediction of biomolecular interactions with AlphaFold 3. Nature. 2024:1–3.

29. DeLano WL. Pymol: An open-source molecular graphics tool. CCP4 Newsl Protein Crystallogr. 2002;40(1):82–92.

30. Trott O, Olson AJ. AutoDock Vina: improving the speed and accuracy of docking with a new scoring function, efficient optimization, and multithreading. J Comput Chem. 2010;31(2):455–461.

31. Van DSD, Lindahl E, Hess B, Groenhof G, Mark AE, Berendsen HJ. GROMACS: fast, flexible, and free. J Comput Chem. 2005;26(16):1701.

32. Genheden S, Ryde U. The MM/PBSA and MM/GBSA methods to estimate ligand-binding affinities. Expert Opin Drug. 2015;10(5):449–461.

33. Chen X, Stafford DW, Tie JK. Assessment of gamma-glutamyl carboxylase activity in its native milieu. Methods Enzymol. 2024;708:207–236.

34. Hao Z, Jin DY, Stafford DW, Tie JK. Vitamin K-dependent carboxylation of coagulation factors: insights from a cell-based functional study. Haematologica. 2020;105(8):2164–2173.

35. Soute BA, Ulrich MM, Vermeer C. Vitamin K-dependent carboxylase: increased efficiency of the carboxylation reaction. Thromb Haemost. 1987;57(1):77–81.

36. Napolitano M, Mariani G, Lapecorella M. Hereditary combined deficiency of the vitamin K-dependent clotting factors. Orphanet J Rare Dis. 2010;5:21.

37. Johnsen JM, Fletcher SN, Huston H, et al. Novel approach to genetic analysis and results in 3000 hemophilia patients enrolled in the My Life, Our Future initiative. Blood Adv. 2017;1(13):824–834.

38. Tokunaga F, Wakabayashi S, Koide T. Warfarin causes the degradation of protein C precursor in the endoplasmic reticulum. Biochemistry. 1995;34(4):1163–1170.

39. Parker CH, Morgan CR, Rand KD, Engen JR, Jorgenson JW, Stafford DW. A conformational investigation of propeptide binding to the integral membrane protein gamma-glutamyl carboxylase using nanodisc hydrogen exchange mass spectrometry. Biochemistry. 2014;53(9):1511–1520.

40. Morris DP, Stevens RD, Wright DJ, Stafford DW. Processive post-translational modification. Vitamin K-dependent carboxylation of a peptide substrate. J Biol Chem. 1995;270(51):30491–30498.

41. Presnell SR, Tripathy A, Lentz BR, Jin DY, Stafford DW. A novel fluorescence assay to study propeptide interaction with gamma-glutamyl carboxylase. Biochemistry. 2001;40(39):11723–11733.

42. Lin PJ, Straight DL, Stafford DW. Binding of the factor IX gamma-carboxyglutamic acid domain to the vitamin K-dependent gamma-glutamyl carboxylase active site induces an allosteric effect that may ensure processive carboxylation and regulate the release of carboxylated product. J Biol Chem. 2004;279(8):6560–6566.

43. Rishavy MA, Hallgren KW, Yakubenko AV, Shtofman RL, Runge KW, Berkner KL. Bronsted analysis reveals Lys218 as the carboxylase active site base that deprotonates vitamin K hydroquinone to initiate vitamin K-dependent protein carboxylation. Biochemistry. 2006;45(44):13239–13248.

44. Rost S, Geisen C, Fregin A, Seifried E, Muller CR, Oldenburg J. Founder mutation Arg485Pro led to recurrent compound heterozygous GGCX genotypes in two German patients with VKCFD type 1. Blood Coagul Fibrinolysis. 2006;17(6):503–507.

45. Lunghi B, Redaelli R, Caimi TM, Corno AR, Bernardi F, Marchetti G. Novel phenotype and gamma-glutamyl carboxylase mutations in combined deficiency of vitamin K-dependent coagulation factors. Haemophilia. 2011;17(5):822–824.

46. Darghouth D, Hallgren KW, Shtofman RL, et al. Compound heterozygosity of novel missense mutations in the gamma-glutamyl-carboxylase gene causes hereditary combined vitamin K–dependent coagulation factor deficiency. Blood. 2006;108(6):1925–1931.

47. Lin PJ, Jin DY, Tie JK, Presnell SR, Straight DL, Stafford DW. The putative vitamin K-dependent gamma-glutamyl carboxylase internal propeptide appears to be the propeptide binding site. J Biol Chem. 2002;277(32):28584–28591.

48. Pudota BN, Hommema EL, Hallgren KW, McNally BA, Lee S, Berkner KL. Identification of sequences within the gamma-carboxylase that represent a novel contact site with vitamin K-dependent proteins and that are required for activity. J Biol Chem. 2001;276(50):46878–46886.

49. Mutucumarana VP, Acher F, Straight DL, Jin DY, Stafford DW. A conserved region of human vitamin K-dependent carboxylase between residues 393 and 404 is important for its interaction with the glutamate substrate. J Biol Chem. 2003;278(47):46488–46493.

50. Gaudry M, Bory S, Dubois J, Azerad R, Marquet A. Vitamin K dependent carboxylation: study of diastereoisomeric gamma-methylglutamic acid containing peptidic substrates. Biochem Biophys Res Commun. 1983;113(2):454–461.

51. Rishavy MA, Berkner KL. Vitamin K oxygenation, glutamate carboxylation, and processivity: defining the three critical facets of catalysis by the vitamin K-dependent carboxylase. Adv Nutr. 2012;3(2):135–148.

52. Shen G, Gao M, Cao Q, Li W. The Molecular Basis of FIX Deficiency in Hemophilia B. Int J Mol Sci. 2022;23(5).

53. Rishavy MA, Berkner KL. Insight into the Coupling Mechanism of the Vitamin K-Dependent Carboxylase: Mutation of Histidine 160 Disrupts Glutamic Acid Carbanion Formation and Efficient Coupling of Vitamin K Epoxidation to Glutamic Acid Carboxylation. Biochemistry. 2008;47(37):9836–9846.

54. Dowd P, Hershline R, Ham SW, Naganathan S. Vitamin K and energy transduction: a base strength amplification mechanism. Science. 1995;269(5231):1684–1691.

55. Tie J, Wu SM, Jin D, Nicchitta CV, Stafford DW. A topological study of the human gamma-glutamyl carboxylase. Blood. 2000;96(3):973–978.

56. Carlisle TL, Suttie JW. Vitamin K dependent carboxylase: subcellular location of the carboxylase and enzymes involved in vitamin K metabolism in rat liver. Biochemistry. 1980;19(6):1161–1167.

